# Optogenetic reactivation of prefrontal social memory trace mimics social buffering of fear

**DOI:** 10.1101/752386

**Authors:** Vanessa A. Gutzeit, Kylia Ahuna, Tabia L. Santos, Ashley M. Cunningham, Meghin Sadsad Rooney, Christine A. Denny, Zoe R. Donaldson

## Abstract

Social buffering occurs when the presence of a companion attenuates the physiological and/or behavioral effects of a stressful or fear-provoking event. It represents a way in which social interactions can immediately and potently modulate behavior. As such, social buffering is one mechanism by which strong social support increases resilience to mental illness. While the behavioral and neuroendocrine impacts of social buffering are well studied in multiple species, including humans, the neuronal bases of this behavioral phenomenon remain largely unexplored. Previous work has shown that the infralimbic prefrontal cortex (IL-PFC) is important for processing social information and, in separate studies, for modulating fear and anxiety. Thus, we hypothesized that socially-active cells within the IL-PFC may integrate social information to modulate fear responsivity. To test this hypothesis, we employed social buffering paradigms in male and female mice. Similar to prior studies in rats, we found that the presence of a cagemate reduced freezing in fear and anxiety-provoking contexts. In accordance with previous work, we demonstrated that interaction with a novel or familiar conspecific induces activity in the IL-PFC as evidenced by increased immediate early gene (IEG) expression. We then utilized an activity-dependent tagging murine line, the ArcCreER^T2^ mice, to express channelrhodopsin (ChR2) in neurons active during the social encoding of a new cagemate. We found that optogenetic reactivation of these socially-active neuronal ensembles phenocopied the effects of cagemate presence in male and female mice in learned and innate fear contexts without being inherently rewarding or altering locomotion. These data suggest that a social neuronal ensemble within the IL-PFC may contribute to social buffering of fear. These neurons may represent a novel therapeutic target for fear and anxiety disorders.

## Introduction

Strong social support in humans is intricately intertwined with our health and well-being [1–3]. The presence of strong social relationships decreases one’s likelihood to develop mental illness and enhances our ability to recover from heart attacks, cancer, and other illnesses [4–7]. In contrast, social isolation has been extensively linked with deleterious health outcomes, ranging from increased rates of coronary heart disease and stroke to neurological and mental health disorders [8]. Adults who are socially isolated or unhappy about their social relationships are at increased risk of premature mortality comparable to the risk posed by obesity or lack of physical activity [2], a finding that has also been observed among other non-human primates that form complex social relationships [9]. Together these epidemiological findings suggest that healthy social relationships have multiple beneficial functions while isolation or abusive relationships are deleterious.

One proposed mechanism by which positive social interactions modulate health is by reducing the behavioral, physiological, and neural response to stress or threat, a phenomenon referred to as social buffering [10,11]. Social buffering is highly conserved across taxa [12,13], and has been documented in rats [14,15], zebrafish [16], goats [17], pigs [18], non-human primates [19–23], and humans [24]. In humans, social buffering consistently attenuates behavioral fear responses and anxiety levels in both experimental and self-reported contexts [25]. Squirrel monkeys exhibit ameliorated stress responses after exposure to a fearful stimulus (a snake) if social companions were present during exposure [26]. In rats, stress responses to fearful stimuli were reduced in the presence of a same-sex partner [27,28], and in mice, the presence of cage-mates reduces anxiolytic responses to novel and aversive stimuli [29].

Given the highly conserved nature of social buffering, its underlying neural mechanisms are hypothesized to be shared across taxa [12,30]. In particular, the PFC modulates social behavior [31–33], fear responses [34,35], and resilience to stressors [36–38]. Critically, inhibition of the PFC with the GABA agonist, muscimol, blocks the effects of social familiarity in reducing anxiety-like behaviors (Lungwitz et al., 2014), suggesting a potentially direct role for the PFC in social buffering of fear and/or anxiety. However, the PFC is a heterogeneous brain structure, and the specific neuronal populations that underlie social buffering remain largely unknown.

Previous work suggests that neuronal ensembles consisting of sparse, interspersed cells mediate distinct behaviors [39]. These neuronal ensembles are often referred to as memory traces or engrams [40,41]. Social engrams in the hippocampus modulate social recognition memory [42], and memory traces in the ventromedial hypothalamus encode social fear [43], suggesting that socially-active neurons in different brain regions encode different aspects of social behavior and social experience. Thus, given the role of the PFC in social buffering and fear learning, we asked whether social memory traces within this brain region integrate social experience and were sufficient to modulate fear and anxiety. We focused on the IL-PFC because of its known role in fear learning and extinction [35,44].

Here, we first validated behavioral metrics of social buffering of fear in mice using a paradigm similar to that previously employed in rats [15]. We subsequently showed that social interaction with a novel or familiar cagemate results in increased expression of the IEG, c-fos, in the IL-PFC. Then, in order to gain genetic access to social memory traces in the IL-PFC, we used ArcCreER^T2^ mice. This mouse line takes advantage of the IEG promoter *Arc* to indelibly label active neuronal populations in a temporally-specific manner [45]. Using the ArcCreER^T2^ mice, we tagged neurons that were active during the introduction of a new cagemate with ChR2-enhanced yellow fluorescent protein (eYFP). Subsequent optogenetic activation of the labeled social memory traces reduced freezing in innate and learned fear tasks, mimicking the behavioral effects of social buffering without altering locomotion or reward/aversion. These data suggest that social memory traces in the IL-PFC may mediate the effects of social buffering. Given that social buffering remains one of the most potent natural regulators of fear and anxiety, targeting this cell population represents a novel therapeutic opportunity.

## Materials and Methods

### Animals and housing conditions

All mice were bred in-house using a mixed background consisting mostly of 129S6/SvEv with a small amount of C57Bl/6J. For all experiments, mice were generated by crossing ArcCreER^T2^ (+) mice [45] with Ai32 heterozygous or homozygous mice (Rosa^ChR2-eYFP^ or Rosa^ChR2-eYFP/ChR2-eYFP^) obtained from Jackson Lab [46]. These mice contain a STOP-floxed ChR2-eYFP integrated at the ROSA locus, resulting in ArcCreER^T2^::Rosa^ChR2-eYFP^, ArcCreER^T2^, Rosa^ChR2-eYFP^ and wildtype pups. This ensured that all animals had the same genetic background across experiments. Genotyping was performed as previously described [45–47]). Male and female mice began experimental testing between post-natal day (PND) 55 and PND 95 (ferrule implantation in optogenetic experiments occurred up to 14 days prior to this).

Social companion animals consisted of C57Bl/6J mice from Jackson Laboratories that were ovariectomized in-house upon arrival, allowed to recover for at least 2-weeks, and used until they reached one year of age. The same animals were used as companions across multiple experiments.

Mice were housed 4 to 5 per cage until experiments began. All cages containing animals in active experiments were changed by experimenters and all cage changes occurred at least 48 hours prior to behavioral testing. Mice were maintained on a 12:12 hour light:dark cycle (06:00-18:00 lights on) at 21 – 25°C with *ad libitum* food and water. All behavioral experiments took place during the light cycle. All animal procedures were approved by the New York State Psychiatric Institute’s and University of Colorado’s Institutional Animal Care and Use Committees (IACUCs).

### Ovariectomy surgery

In order to avoid the confounds of sexual receptivity and pregnancy, female companion mice were ovariectomized [48]. Mice were anesthetized and maintained on 1 – 3% isoflurane and depth of anesthesia was monitored via foot pinch and breathing. Paralube was applied to the eyes to avoid drying and core body temperature was monitored and adjusted via a rectal thermometer and a heated pad placed underneath the animal. Hair was shaved at the incision site and the underlying skin was disinfected with betadine and 70% ethanol (EtOH). A single incision was made in the midline of the back to allow access to the body cavity. the incision was pulled to one side until aligned above the ovary. A small incision was made in the body wall, the ovary was pulled through, removed, and the edges of the uterus were cauterized. The internal body wall was closed using an absorbable suture, the skin was pulled to the other side, and the procedure was repeated. The external incision was closed with staples, which were removed 7 days later. Triple antibiotic ointment and lidocaine were places on the closed wound, and mice received a single dose of Meloxicam SR (2 mg/kg) for analgesia.

### Social buffering of fear responses

To assess the effects of conspecific presence on freezing in fear-provoking environments, we conducted a series of behavioral tests. Experimental animals were singly housed in fresh cages for 48 hours, and then provided with an ovariectomized female cagemate for the remainder of the study. In the first series of behavioral tests, mice were fear conditioned 5 days after cagemate introduction. Freezing was measured when the test animals were re-exposed to the context with or without their cagemate 2 days later. Two days later, freezing levels were measured when test animals were exposed to a brightly lit, novel arena with or without their cagemate. In a separate, independent experiment we also measured freezing when animals were exposed to a lemon scent that was present during fear conditioning (FC). For all behavioral tests, mice were habituated to the room at least 30 minutes before testing. Details of behavioral testing are provided below (Fig 1A).

**Figure 1.**
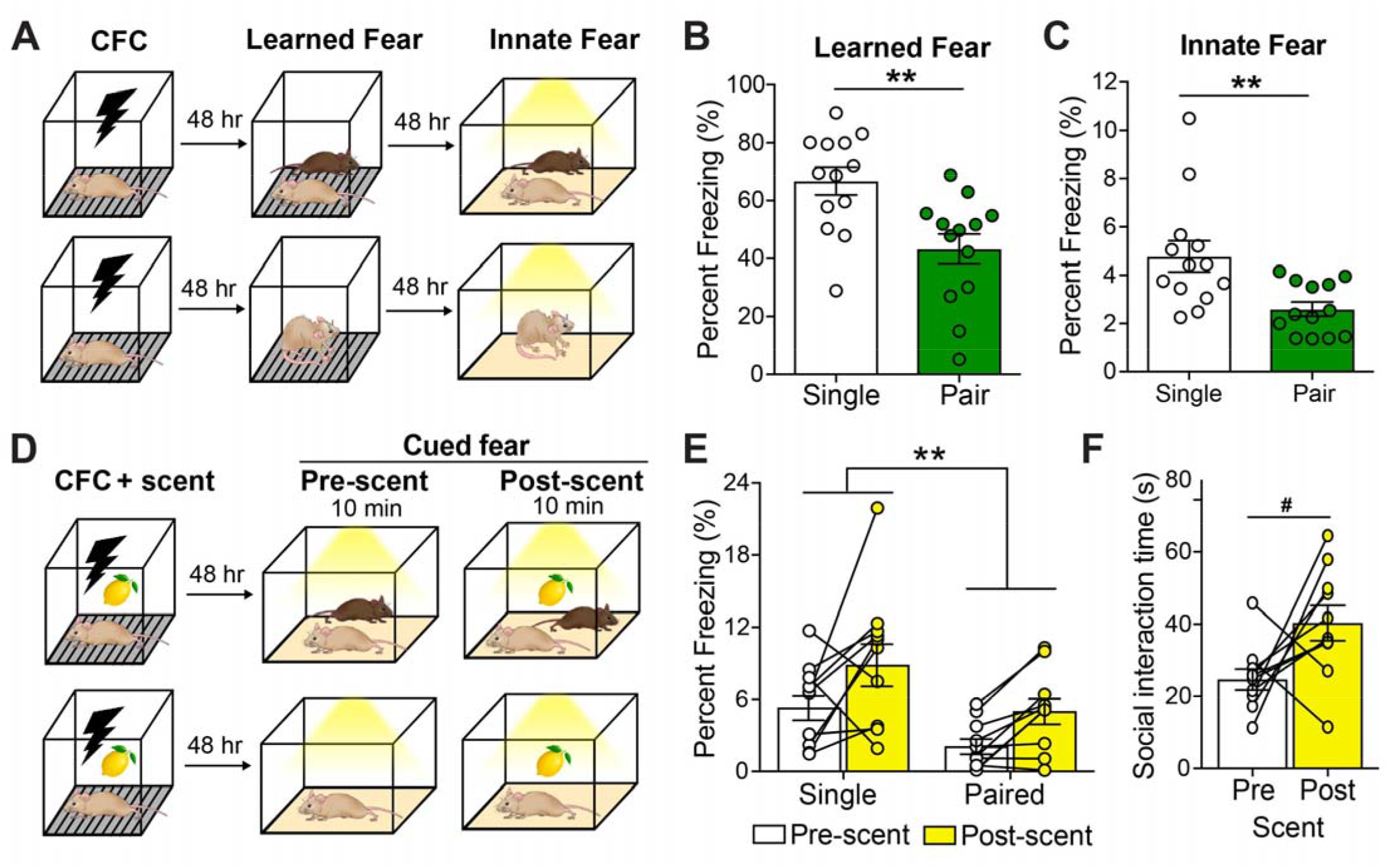
Cagemate presence reduces freezing in multiple fear-provoking tests. (A) Timeline and behavioral schematic of testing with (top) or without (bottom) a cagemate. Learned fear was assessed by measuring freezing upon re-exposure to a context previously paired with an aversive shock while innate fear was assessed in an anxiogenic novel environment. (B) Mice re-exposed to an aversive context with their cagemate showed lower levels of freezing when compared to mice tested alone. (C) Similar to learned fear, mice froze less in an innate fear context when tested with their cagemate present compared to mice tested alone. (D) Timeline and behavioral schematic showing aversive cue exposure with (top) or without (bottom) a cagemate present. After receiving a shock with a lemon scent present, mice were exposed to a novel context for 5 minutes and then the lemon scent (cue) was added to the arena for another 5 minutes. The lemon scent was the only shock-associated cue present in the test. (E) All mice increased their freezing levels after presentation of the lemon scent, but cagemate presence reduce freezing before and after scent presentation relative to animals tested alone. (F) Paired mice showed a trend towards increased interaction with their cagemate after the scent (fear cue) was added to the chamber. Error bars represent ± SEM. * p < 0.05; ** p < 0.01, *** p < 0.001. # p = 0.055 CFC, contextual fear conditioning; hr, hour.

#### Learned fear

Contextual FC was performed as previously reported [45,49]. FC chambers were from Coulbourn Instruments with the internal dimensions of 7 × 7 × 12 in. The chambers had clear plastic front and back walls, stainless-steel walls on each side, and stainless-steel bars on the floor. A house light was mounted directly above the chamber. Each chamber was located inside a larger, insulated plastic cabinet that provided protection from outside light and noise. Each cabinet contained a ventilation fan that was operated during the sessions. A paper towel dabbed with lemon solution was placed under the stainless-steel bars. The scent was refreshed for each animal. Mice were held outside the experimental room in their home cages prior to testing and transported to the conditioning apparatus in standard mouse cages. FC chambers were cleaned with 70% EtOH between each run. During training, mice were placed in the chamber. They received a 2 sec 0.75 mA shock at 180, 240, and 300 seconds after introduction into the chamber. The animal was removed 15 seconds after the last shock. Fear memory retrieval was assessed by returning the mice to the fear conditioning chamber 48 hours later for 5 minutes with or without their cagemate (no shock was delivered during this phase). Freezing of the test animal was hand-scored in a blinded fashion. Freezing levels were compared using a two-way ANOVA with Sex and Cagemate Presence (treatment) as between-subject factors.

#### Innate fear

To assess innate fear, mice were exposed with or without their cagemate to a novel environment for ten minutes. The testing occurred in a different room than fear conditioning. The novel environment consisted of square chambers (16 × 16 in) with metal floors, no bedding, and clear walls. All the lights in the room were turned on to maximum output (lux = 200). Chambers were cleaned between each test with PDI™ Super Sani-Cloth™ Germicidal Disposable Wipes. Mice were kept inside the room during testing and were transferred from the home cage to the center of the chamber by the experimenter. Freezing levels were compared using a two-way ANOVA with Sex and Cagemate Presence (treatment) as between-subject factors.

#### Cued fear

Mice were FC as described above, where a paper towel dabbed with lemon solution was placed under the stainless-steel bars, and the scent was refreshed for each animal. Two days after this test, mice were placed in a novel environment of a rectangular white chamber (12 × 24 in) with or without their partner. These cages contained different bedding from their home cage. All lights in the room were turned on to maximum output (lux = 200). Mice were kept inside the room during testing and were transferred from the home cage to a randomized corner of the chamber by the experimenter, and behavior was recorded via a camera above the chamber. A 5-minute baseline was taken in the chamber (pre-scent), and then a swab of the lemon scent was added to the chamber, and 5 additional minutes were recorded (post-scent). Bedding was changed between each mouse, and freezing levels and time in center were hand scored by an experimenter blinded to condition. Freezing levels were compared using a two way ANOVA with Sex and Cagemate Presence (treatment) as between-subject factors.

### IL-PFC c-fos levels following social interaction

#### c-fos induction

Animals were transferred to a fresh individual cages and housed for 48 hours with an ovariectomized female. To examine patterns of c-fos induction in response to novel or familiar mouse or a novel object, the ovariectomized female was removed from the cage for 1 hour, and then either a novel object (50 ml conical tube), novel ovariectomized female, or the ovariectomized cagemate was placed in the test animal’s home cage. The stimulus animal/object was removed after 50 minutes, and test animals were perfused and brain tissue was collected 60 minutes after initial stimulus introduction.

#### Immunolabeling of c-fos

Mice were deeply anesthetized with (*R*,*S*)-ketamine (100 mg/kg) and xylazine (10 mg/kg) and were transcardially perfused with 0.1M phosphate buffered saline (1X PBS) followed by 4% paraformaldehyde (PFA)/1X PBS. The brains were post-fixed overnight in 4% PFA at 4°C, then transferred to 30% sucrose/1X PBS until the brains dropped to the bottom of the tubes. Serial coronal sections (30 μm) were cut, and floating sections were used for all labeling procedures.

Sections were rinsed three times in 1X PBS and blocked for 2 hours at room temperature in 1X PBS with 0.3% Triton-X (0.3% PBST) with 10% normal donkey serum (NDS). Sections were incubated in primary antibody in 0.3% PBST/ 3% NDS (rabbit anti-c-fos, Synaptic Systems, 226003, 1:5000) for 72 hours at 4°C. Sections were washed three times with 1X PBS and blocked in 0.3%PBST/2%NDS for 1 hour at room temperature. Sections were incubated in Cy3-conjugated donkey anti-rabbit (Jackson Immunoresearch, 711-165-152, 1:500) secondary antibodies for 2 hours in 1X PBS with 0.25% Tween. Sections were washed three times in 1X PBS and mounted on slides. Once dry, slides were coverslipped using Prolong Diamond reagent (Life Technologies P36965), allowed to dry and stored at 4°C until imaging.

#### Microscopy and cell counts

Three consecutive tissue sections separated by 100 μm were used for counting c-fos^+^ cells. Images were acquired using an Olympus VS120 slide scanning microscope. For each animal, equivalent sections of the PFC that contained the infralimbic cortex were chosen for cell counting. The Allen Brain Atlas was used as a reference, and the same overlay was used to define the region for counting c-fos^+^ cells in each animal (Fig 2A). c-fos^+^ cells were manually counted, and cell counts were averaged across the left and right hemispheres of 3 matched sections per mouse.

**Figure 2.**
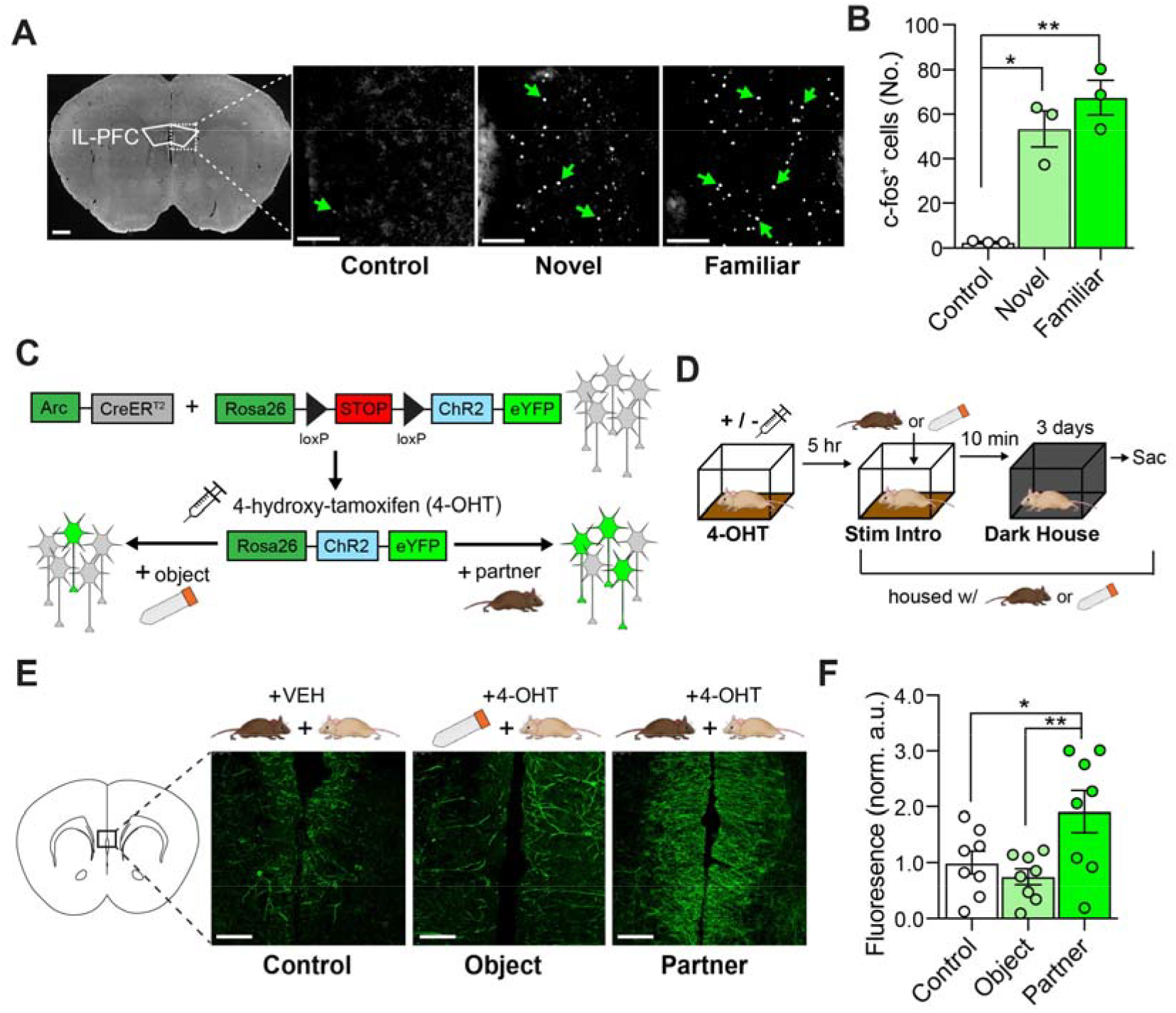
The IL-PFC exhibits robust IEG induction and IEG-mediated cellular tagging following social interaction. (A) Representative images of c-fos induction following exposure to a novel or familiar mouse, compared to controls exposed to a novel object (conical tube). Scale bars left = 10 mm, right = 50 µm. (B) Quantification of c-fos^+^ cells in mice exposed to a novel object (control; 2.5 ± 1.258, n = 3), a novel mouse (59.83 ± 14.34, n = 6), or a familiar mouse (97.63 ± 18.42, n = 4). Exposure to a novel or familiar mouse results in significant induction of c-fos^+^ cells relative to object-exposed controls, but there were no differences between socially exposed groups (p = 0.98). (C) ArcCreER^T2^ × ROSA26-STOP-floxed-ChR2-eYFP mice were used to indelibly label socially-active neurons (social memory traces) with ChR2-eYFP. Briefly, CreER^T2^ is under the control of the Arc promoter. Upon administration of 4-OHT, CreER^T2^ localizes to the nucleus and recombines loxP sites in the floxed-STOP-ChR2-eYFP transgene, resulting in indelible expression of ChR2-eYFP in Arc^+^ cells. Labelling ends when 4-OHT is metabolized and excreted. (D) Timeline for labeling a social memory trace in the IL-PFC. Mice received 4-OHT or vehicle, and 5 hours later were exposed to a new cagemate or novel object. (E) Representative images of immunolabeling of ChR2-eYFP^+^ cells in the IL-PFC in animals receiving 4-OHT or vehicle prior to exposure to a novel mouse or a novel object. Scale bars = 50 µm (F) Quantification of eYFP in the IL-PFC, where robust labeling was observed in animals receiving 4-OHT and exposed to a novel mouse compared to an object or vehicle controls. Error bars represent ± SEM. * p < 0.05; ** p < 0.01, *** p < 0.001. Veh, vehicle; 4-OHT, 4-hydroxytamoxifen; stim, stimulus; hr, hour; min, minutes; sac, sacrifice; norm, normalized.

### Labeling social memory traces in the IL-PFC

#### ArcCreER^T2^-mediated cell labeling

Recombination was induced with 4-hydroxytamoxifen (4-OHT, Sigma-Aldrich). 4-OHT was dissolved by sonication in 10% EtOH/90% corn oil as a concentration of 10 mg/ml. All animals were singly housed for 2 days. On the morning of day 3, animals received a single intraperitoneal (i.p.) injection of 0.15 ml 4-OHT or vehicle (VEH) (10% EtOH/90% corn oil) (Fig 2C) [47]. Five hours after injection, animals were introduced to and subsequently lived with an ovariectomized female or novel object. Cages were left undisturbed with the lights off for 60 hours until the normal light-dark cycle was resumed (Fig 2D). Experimental animals consisted of ArcCreER^T2^::Rosa^ChR2-eYFP^ that received 4-OHT.. Controls were ArcCreER^T2^::Rosa^ChR2-eYFP^ that received vehicle or Rosa^ChR2-eYFP^, ArcCreER^T2^, or wildtype littermates mice that received 4-OHT.

#### Measuring eYFP induction

Brains were collected 5 days after cagemate/object introduction to measure eYFP induction. Animals were perfused and sections were prepared as described above. eYFP signal was immuno-amplified using the labeling methods outlined above but with chicken anti-GFP polyclonal primary antibody (Abcam, ab13970, 1:500) for 24 hours at 4°C and Cy2-conjugated donkey anti-chicken secondary (Jackson Immunoresearch, 703-225-155, 1:500). Because ChR2-eYFP is localized to the membrane, we were able to visualize large arborizations in the mPFC; however, the density of the membrane-bound fluorophore within this region made if challenging to identify individual eYFP+ soma. (Fig 2E). Thus, we quantified relative ChR2-eYFP induction by measuring overall fluorescence in the IL-PFC in matched sections (Fig 2F). We measured fluorescent intensity in ImageJ across 8 images for each condition (VEH, object, and partner) by taking the mean fluorescence of the image and subtracting the background fluorescence from a region of interest with no visible eYFP labeling, and this value was then normalized to the intensity of the control group. As we observed the most visually robust eYFP induction in the posterior portion of the IL-PFC, we focused our analysis there for measuring fluorescent intensity, c-fos counts (see above), and for optogenetic experiments. In contrast to sparsely labeled areas, such as the dentate gyrus [47], dense labeling with membrane-bound eYFP in the IL-PFC made full co-localization of a somatic IEG extremely difficult.

### Optogenetic manipulation of social memory traces

#### Construction of optical fibers

Optical fibers for ferrules were constructed using techniques published by [50]. A 2.5 mm length, 200-μm core with 0.37 numerical aperture (NA) multimode fiber was glued with epoxy and threaded through a 230-μm core zirconia multimode ferrule (ThorLabs, Newton, NJ). The ferrules were polished. Implants were tested for light output, with greater than ~70% light recovery required using a light meter (Thorlabs, Newton, NJ) and acceptable light cone emission used as criteria for use. Subsequently, each acceptable ferrule was numbered and the percent of light recovery was recorded for calibration of light output during behavior experiments. An optical patch cable (Doric lenses, Quebec, QC, Canada) was used to connect the ferrule to a multimode FC ferrule assembly at a 1 × 2 optical rotary joint (Doric lenses, Quebec, QC, Canada). The other end of the rotary joint was connected via a patch cable to a 100 mW 473 nm blue laser (Shanghai Laser & Optics Century Co.) via a non-contact-style laser to fiber coupler (OZ optics, Carp, ON, Canada).

#### Ferrule implantation

Mice were induced and maintained on 1 – 3% isoflurane anesthesia. They were placed in a stereotactic frame (Kopf). The scalp was opened using a midline incision. The skull was scored and a small screw was inserted to help anchor the implanted ferrule. A 1 mm hole was drilled using a 22-gauge drill bit at the site of implantation. A single fiberoptic was implanted into the midline IL-PFC using the following coordinates – AP +1.8mm, DV −2.3mm, ML+0.33 (to avoid hitting the medial artery). Dental cement or Loctite 454 with blackening charcoal powder were used to secure the fiberoptic implant. The incision was closed using VetBond. Lidocaine ointment was applied to the incision, and all animals received injections of carprofen (1 mg/kg) solution for 3 days following surgery. Animals recovered for at least 10 days prior to initiation of experiments.

#### ArcCreER^T2^-mediated cell labeling

Social memory traces were labeled as described above, but all animals (genotypes for experimental and control are outlined above) were paired with ovariectomized female 5 hours after VEH or 4-OHT administration (Fig 3A). Experimental animals continued to live with this female through the duration of the experiment. Behavioral testing was initiated 10 – 14 days after the VEH or 4-OHT injection to ensure robust expression of ChR2-eYFP.

**Figure 3.**
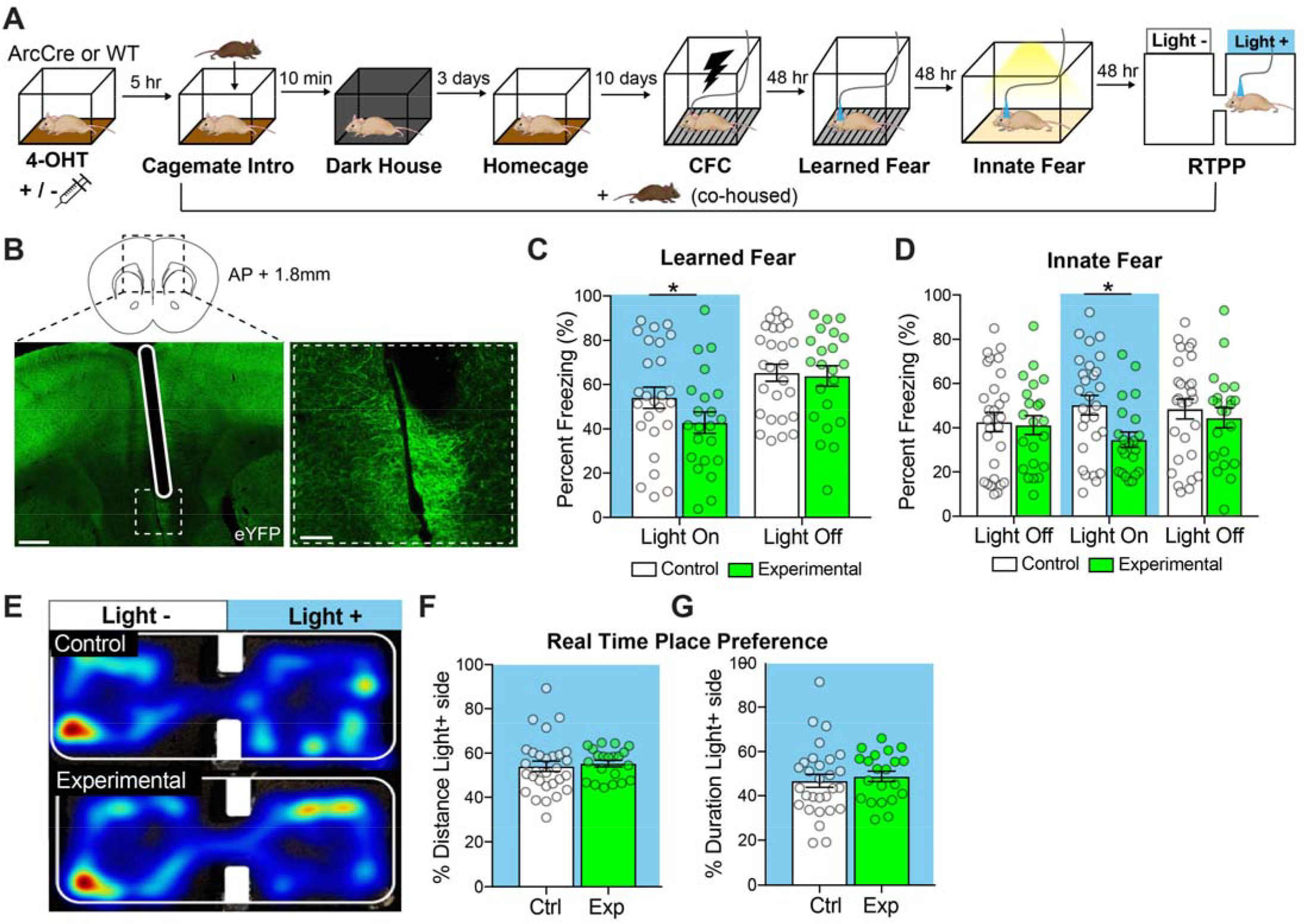
Optogenetic reactivation of socially labeled neuronal ensembles reduces freezing in multiple tests. (A) Behavioral timeline for optogenetic cohorts. (B) Visualization of fiberoptic implanted in the midline of the IL-PFC (below), and magnified image of eFYP socially labeled cells in the IL-PFC (right). Scale bars left = 100 µm, right = 50 µm. (C) Optogenetic reactivation of the social memory trace in the IL-PFC caused a reduction in freezing upon re-exposure to an environment previously paired with a shock. (D) Optogenetic reactivation of socially-labeled ensembles significantly reduced freezing in a novel context during light-on period. (E) Representative heat maps for locomotion during RTPP (top = control, bottom = experimental). Optogenetic stimulation of socially-labeled cells in the IL-PFC was not aversive or rewarding. The percent distance traveled (F) and the duration (G) in the light-on chamber was not significantly different between control and experimental animals. Error bars represent ± SEM. * p < 0.05; ** p < 0.01, *** p < 0.001. 4-OHT, 4-hydroxytamoxifen; CFC, contextual fear conditioning; RTPP, real time place preference; hr, hours; min, minutes; blue = light ON.

#### Optogenetics manipulation

For all optogenetic experiments, the laser was turned on at least 30 minutes before and tested for consistent light output prior to experimentation. The fiberoptic implant was cleaned with an IPA wipe (VWR) prior to attaching the patch cable. Output was normalized by individual implant for each mouse to 10-12 mW. A master8 or custom Arduino interface was used to control light delivery, achieving 5 msec pulses at 10 Hz.

#### Behavioral testing

For all behavioral experiments assessing the effects of activation of the social memory trace, mice lived with their cagemate but were tested alone. Experimental mice underwent 3 tests in the following order: conditioned fear assessment (learned fear), exposure to a novel environment (innate fear), and real time place preference (RTPP) (Fig 3A).

#### Learned fear

Mice underwent FC training as outlined above. Freezing as a measure of fear memory was assessed by returning the animal to the training context 48 hours later. Test animals were attached to the laser and introduced to the chamber with the laser one. After 3 minutes, the laser was turned off, enabling a within-animal measure of freezing. Freezing levels were hand scored in a blinded fashion, and were analyzed using a repeated measures ANOVA with Laser ON/OFF as a within-subject factor and Group (control or experimental) as a between subject factor.

#### Innate fear

Freezing in a novel context was assessed as described above. Animals were introduced to the novel environment with the fiberoptic attached and allowed to explore for three minutes with the laser off, light was then applied for three minutes, and was removed for three minutes, resulting in a total test time of nine minutes. Freezing levels were hand scored by an experimenter blinded to condition. Freezing levels were analyzed using a repeated measures ANOVA with Laser On/Off as a within-subject factor and Group (experimental or control) as a between subject factor.

#### Real Time Place Preference

In order to determine whether optogenetic activation of the social memory trace had rewarding or aversive properties or affected locomotion, mice were tested in a RTPP task in which laser stimulation was delivered on one side of a two chambered apparatus [51]. The arena (two 8.25 × 9.25 in chambers connected by a 4-inch opening) contained bedding and was placed on the floor in the same room as the novel context, with one center light shining on the apparatus). The test animal’s location was tracked using Ethovision XT 10 (Noldus), which interfaced with the laser to deliver continuous 5 msec, 10 Hz light as long as the animal was on the stimulation-paired side. Sessions ran for 20 minutes, and stimulation side was alternated between animals. Time and distance traveled on each side of the apparatus were recorded in 5-minute bins. Group differences for locomotion and for percentage of time or distance traveled on the light-paired side were assessed using a repeated measures ANOVA with Time as a within-subject factor and Group as a between-subject factor.

### Data Analysis

All data are presented as mean +/− SEM. Data analysis was conducted using Statistical Package for the Social Sciences (SPSS) version 24 IBM software. Specific statistical tests used are indicated above, and detailed results of statistical analyses and group sizes are available in **Supplemental Table 1**.

## Results

### Social buffering occurs in multiple fear-provoking contexts

We sought to develop methods to assess social buffering across a range of tests that varied in the extent to which they elicited freezing. In the most intense of these tests, mice were re-exposed to a context in which they had previously received aversive shocks (Fig 1A). Mice that were placed in this context with their cagemate exhibited lower levels of freezing than mice that were placed in this context alone (Fig 1B). Specifically, Cagemate Presence had a significant main effect on freezing levels (p < 0.001), as did Sex of the experimental animal (p = 0.007; **Fig S1A, B**), but there was no interaction between Cagemate Presence and Sex (p = 0.294). We also exposed the same subjects to a separate context containing anxiogenic environmental cues (bright lights and a solid metal floor without bedding). In order to avoid generalization, this was carried out by a different experimenter in a separate room. Mice who were introduced with their cagemate present exhibited decreased freezing compared with the mice placed in the apparatus alone (Fig 1C). Specifically, Cagemate Presence had a significant main effect on freezing levels (p = 0.005), but there was no effect of Sex of the experimental animal (p = 0.504) or interaction between Cagemate Presence and Sex (p = 0.775) (**Fig S1A, B**). These two tests produced freezing levels that differed by an order of magnitude, suggesting substantial differences in the amount of fear elicited by each test. However, cagemate presence had a significant effect on freezing levels in both tests, suggesting that social buffering effects are evident across a wide range of fear provoking/anxiogenic environments.

In a separate cohort, we also examined the role of cagemate presence on cued olfactory fear responses. After being trained to associate a lemon scent with an aversive shock, the test animal was placed in a novel context with or without their cagemate. A comparison of freezing levels before (pre-scent) and after (post-scent) re-exposure to the lemon scent (cue) (Fig 1D) revealed an increase in freezing following scent introduction (Fig 1E; p = 0.005). However, freezing levels were significantly decreased both pre- and post-scent if the animal was placed in the context with their cagemate (Fig 1E; p = 0.02). When separated by sex, this social buffering effect was significant in males (**Fig S1E**; p = 0.01) but not in females (**Fig S1F**; p = 0.5). However, this may reflect the small number of female mice available for this particular test. In this test, freezing levels before cue introduction were comparable to those observed in the novel environment and increased to an intermediate level following scent application. Finally, among mice that had their cagemate present during testing, the time spent interacting with the cagemate tended to increase following scent addition (Fig 1F; p = 0.055).

Taken together, these results indicate that having a cagemate present blunts freezing in a range of fear-provoking situations, providing experimental evidence that cagemate presence produces a robust social buffering effect. In addition, both males and females are sensitive to cagemate presence, although to a different extent in different tests.

### IL-PFC exhibits robust induction of c-fos following social but not object interaction

Having demonstrated that cagemate presence decreases a behavioral metric of fear, and based on the known role of the IL-PFC in fear modulation, we next asked whether IEG expression was upregulated in the IL-PFC following social interaction. Specifically, we compared c-fos expression in the IL-PFC in mice exposed to a familiar ovariectomized female (familiar), novel ovariectomized female (novel), or a novel object (control) (Fig 2A). We found that there were significantly more c-fos^+^ cells in the IL-PFC of mice exposed to a novel (p = 0.03) or familiar (p = 0.007) mouse as compared with controls (Fisher’s multiple comparisons test). There was no difference in number of c-fos^+^ cells found between the novel and familiar exposure conditions (Fig 2B; p = 0.14). Thus, these data indicate that social interaction substantially increases IEG expression in the IL-PFC.

### Optogenetic activation of social memory traces in IL-PFC reduces freezing

In order to identify and manipulate social memory traces, we used the ArcCreER^T2^ X Ai32 mice to express ChR2-eYFP in socially-active neurons (Fig 2C-D). We observed a greater amount of eYFP expression in the IL-PFC of mice paired with an ovariectomized female as compared to controls (Fig 2E-F; p < 0.05), mirroring socially-induced IEG expression (Fig 2B).

To test whether activation of the social memory trace in the IL-PFC modulates freezing levels, we optogenetically stimulated ChR2-eYFP^+^ tagged neurons in the IL-PFC while mice experienced different fear-provoking environments. The experimental timeline is summarized in Figure 3A. Light-induced activation of the social memory traces reduced freezing when animals were placed in a context in which they had previously been shocked (Fig 3C, **S2A**, Light X Time: p = 0.036). In addition, experimental animals exhibited a greater increase in freezing in the minute following the light being turned off as compared to controls (p = 0.04; **Fig S2b**). Light-induced activation of the same social memory trace also reduced freezing in a novel arena, which returned to control levels when the laser was turned off (Fig 3D, **S2B**; Light X Time: p = 0.01). The effects of memory trace activation were strongest immediately following light onset; control animals did not change their freezing (p = 0.640), but experimental animals decreased their freezing by nearly 20% (p = 0.005) (**Fig. S2B**). Notably, freezing levels in the novel environment were greater in our optogenetic experiments than in prior behavioral tests (e.g. Fig 1C). All animals had the fiberoptic attached to the ferrule during FC. Thus, the fiber-optic cable may have served as an aversive cue and contributed to the elevated freezing levels in the novel context. For all analyses, males and females were combined as there was no main effect or interacting effects of Sex. This indicates that reactivation of socially-active neurons in the IL-PFC mimics the effects of social buffering.

### Reactivation of social neuronal ensembles is not rewarding or aversive and does not alter locomotion

Using a real time place preference (RTPP) test, we found that stimulation of the social memory trace was not aversive or rewarding and did not affect locomotion. Specifically, stimulation of the social memory trace did not alter the amount of time that the mice spent in the light ON chamber compared with the light OFF chamber (Fig 3E, 3G, **S2D**, p = 0.5). Similarly, there was no difference in locomotion when the animals were in the light ON chamber compared with the light OFF chamber (Fig 3F, **S2C**, p = 0.7). For all analyses, males and females were combined as there was no main or interacting effect of sex. Thus the differences in freezing observed during optogenetic manipulation are not attributable to induction of locomotor or reward-related processes.

## Discussion

The buffering of fear responses by conspecific presence is a broadly observed and highly conserved phenomenon. We have demonstrated that companion presence can attenuate freezing in mice and that this effect can be recapitulated in the absence of a conspecific by optogenetically reactivating socially-active neurons in the IL-PFC. Further, we demonstrated that this is effect generalizes across multiple fear-provoking situations that vary in intensity. This suggests that a social memory traces within the IL-PFC may exert top-down control to modulate the behavioral expression of fear and that this is not context- or cue-specific.

Multiple lines of evidence indicate that the IL-PFC processes social information and modulates fear responses, although few studies have examined the potential convergence of these functions. The IL-PFC exhibits robust neuronal activity during social interaction [31–33], and we confirmed that interaction with either a novel or familiar conspecific results in increased c-fos^+^ expression within this region [52]. Likewise, the IL-PFC is required for extinction of fear [44,53,54]. Some studies directly or indirectly suggest a role for this region in social buffering. Specifically, inhibition of the IL-PFC via muscimol infusion acutely blocks the effects of social familiarity in reducing anxiety-like behaviors [55] and lesions of this region abolish the resiliency-promoting effects of environmental enrichment [56,57]. The work presented here suggests that a specific subset of socially-active neurons (the social memory trace) may mediate social buffering, further contributing to our understanding of how social processing and fear learning may be integrated within this brain region.

The prelimbic (PL)- and IL-PFC may be ideally suited to integrate multiple information streams. For instance, recent work revealed a specialized subset of social place cells in the PL-PFC that integrate social and spatial information [32]. The social memory trace in the IL-PFC may similarly integrate social memory and fear learning (or safety cues). But how does this fit into a distributed social memory circuit? Rodents rely on olfactory information to recognize other individuals, and olfactory inputs are required for social buffering in rats [11,58]. Olfactory-based identity information eventually contributes to social memory engram in the ventral CA1 (vCA1) region of the hippocampus (HPC) [59] and projections from this region to the PFC are required for social recognition memory [42,60]. Thus, the social memory trace in the IL-PFC may represent an advanced level of social information processing reliant on social engram projections from vCA1 feeding into mPFC.

One novel aspect of our study is that it suggests that harnessing social memory traces may be useful in ameliorating fear and/or anxiety in diverse tasks and environments. Prior studies have captured a memory linked to a specific fear-provoking event/context [45,61,62]. Subsequent manipulation of these engrams has revealed essential aspects of memory and its plasticity [63–67]. While this may have therapeutic potential for disorders linked to a specific traumatic event, such as PTSD, it provides less promise for addressing generalized anxiety disorder (GAD) or fear disorders. Specifically, activation of IL-PFC social memory traces may be effective in modulating fear and anxiety levels more generally.

While we found that optogenetic reactivation of socially-labeled neurons attenuated fear, there remain a number of questions regarding the identity of these neurons, how they are recruited by social interaction, and how they regulate fear responses within a larger neural circuit. Previous work has shown that the mPFC exerts top-down control of fear and anxiety [68]. Specifically, projections from the IL-PFC to the basolateral amygdala (BLA) are required for fear extinction [35,44]. If social interaction preferentially recruits IL-BLA projecting neurons, this could explain how activation of these cells regulates fear expression. Additionally, oxytocin is required for social buffering, and oxytocin signaling in the prelimbic cortex reduces anxiety [10,14,22,69,70]. One intriguing possibility is that oxytocin signaling directly or via connections from the mPFC recruits the subset of cells in the IL-PFC that are active during social interaction.

Social buffering encompasses a broad range of physiological and behavioral responses [71]. Here, we focused exclusively on the modulation of freezing as a behavioral indication of fear. Previous work has suggested that the physiological and behavioral effects of social buffering may be dissociable; different social exposures can elicit different buffering responses. For instance in rats, conspecific presence while experiencing an anxiogenic environment reduces freezing, while co-housing with a conspecific following fear conditioning during the recovery period attenuates autonomic responses (specifically hypothermia) without altering freezing [28,72]. Other studies have also examined how hypothalamo-pituitary-adrenal axis activity is modulated by conspecific presence [73–75]. Determining whether activation of the social memory trace in the IL-PFC can ameliorate stress responses, in addition to fear, will provide insight into the neural architecture of different facets of social buffering.

In addition to the areas of further investigation outlined above, our approach also has a few limitations. In particular, we focused on freezing as a metric of fear expression, but we did not limit interaction between the test animal and their cagemate during testing. Thus freezing could also be reduced because of the cagemate physically investigating the test animal. These concerns are mitigated because optogenetic activation, where only the test animal was present, resulted in similar decreases in freezing. However, as with all manipulations that measure freezing, it is possible that our interventions did not directly ameliorate fear but instead distracted the subject from a fearful context and/or engaged an exploratory drive. In addition, the scope of our work is limited to gain-of-function optogenetic manipulations, and as such, we cannot make any conclusions about the necessity of socially-active IL-PFC neurons for the effects of social buffering. One potential way to address the requirement of these neurons for social buffering would be to optogenetically inactivate these cells while mice are exposed to fear-provoking environments with a cagemate present. Finally, we tested the effects of only a single type of social stimulus, ovariectomized female cagemates, and it is possible that different social partners could engage different neuronal ensembles, thereby producing different effects on fear levels. The latter has been well documented in rats and to some extent in mice and voles, where the developmental stage, emotional state, and affiliative nature of the companion animal are all important modulators of social buffering [69,76–79].

In summary, social buffering is a powerful regulator of fear and anxiety across species, and this work provides a potential cellular substrate that mediates the effects of companion presence on fear expression. Despite differences in the sensory factors that are important for social buffering in rodents (olfactory/tactile) and humans (visual/tactile), substantial evidence suggests that similar socially-sensitive neural circuits are recruited to reduce fear responses in humans and rodents [30]. Within this circuitry, the PFC is ideally positioned to integrate sensory information and exert top-down social regulation of emotional states [34,80]. As such, understanding the neural dynamics and circuits that underlie social buffering may contribute to novel therapeutics for fear and anxiety disorders.

## Author Contributions and Notes

Z.R.D., C.A.D. designed research, V.A.G., K.A., T.L.S., A.M.C., M.S. performed research, V.A.G., K.A., M.S. analyzed data; and V.A.G. and Z.R.D. wrote the paper.

This article contains supporting information online.

## Acknowledgments

We would like to thank the animal care staff at Columbia University and University of Colorado Boulder. Additional technical assistance was provided by Magdalena Woroniecka, Kelsey Harbart, and Katelyn Gordon. This work was supported by R00 MH102352 and a Young Investigator Grant from the American Foundation for Suicide Prevention (Z.R.D.)

## References

1. DeVries AC, Glasper ER, Detillion CE. Social modulation of stress responses. Physiol Behav. 2003;79:399–407.

2. Holt-Lunstad J, Smith TB, Layton JB. Social Relationships and Mortality Risk: A Meta-analytic Review. PLOS Medicine. 2010;7:e1000316.

3. House JS, Landis KR, Umberson D. Social relationships and health. Science. 1988;241:540–545.

4. Ozbay F, Fitterling H, Charney D, Southwick S. Social support and resilience to stress across the life span: a neurobiologic framework. Curr Psychiatry Rep. 2008;10:304–310.

5. Ozbay F, Johnson DC, Dimoulas E, Morgan CA, Charney D, Southwick S. Social support and resilience to stress: from neurobiology to clinical practice. Psychiatry (Edgmont). 2007;4:35–40.

6. Reblin M, Uchino BN. Social and emotional support and its implication for health. Curr Opin Psychiatry. 2008;21:201–205.

7. Rodríguez-Artalejo F, Guallar-Castillón P, Herrera MC, Otero CM, Chiva MO, Ochoa CC, et al. Social Network as a Predictor of Hospital Readmission and Mortality Among Older Patients With Heart Failure. Journal of Cardiac Failure. 2006;12:621–627.

8. Valtorta NK, Kanaan M, Gilbody S, Ronzi S, Hanratty B. Loneliness and social isolation as risk factors for coronary heart disease and stroke: systematic review and meta-analysis of longitudinal observational studies. Heart. 2016;102:1009–1016.

9. Archie EA, Tung J, Clark M, Altmann J, Alberts SC. Social affiliation matters: both same-sex and opposite-sex relationships predict survival in wild female baboons. Proc Biol Sci. 2014;281.

10. Kikusui T, Winslow JT, Mori Y. Social buffering: relief from stress and anxiety. Philosophical Transactions of the Royal Society of London Series B, Biological Sciences. 2006;361:2215–2228.

11. Kiyokawa Y, Wakabayashi Y, Takeuchi Y, Mori Y. The neural pathway underlying social buffering of conditioned fear responses in male rats. Eur J Neurosci. 2012;36:3429–3437.

12. Beery AK, Kaufer D. Stress, social behavior, and resilience: Insights from rodents. Neurobiology of Stress. 2015;1:116–127.

13. Bovard EW. The effects of social stimuli on the response to stress. Psychol Rev. 1959;66:267–277.

14. Armario A, Luna G, Balasch J. The effect of conspecifics on corticoadrenal response of rats to a novel environment. Behavioral and Neural Biology. 1983;37:332–337.

15. Davitz JR, Mason DJ. Socially facilitated reduction of a fear response in rats. J Comp Physiol Psychol. 1955;48:149–151.

16. Faustino AI, Tacão-Monteiro A, Oliveira RF. Mechanisms of social buffering of fear in zebrafish. Scientific Reports. 2017;7:44329.

17. Lyons DM, Price EO, Moberg GP. Social grouping tendencies and separation-induced distress in juvenile sheep and goats. Dev Psychobiol. 1993;26:251–259.

18. Kanitz E, Hameister T, Tuchscherer M, Tuchscherer A, Puppe B. Social support attenuates the adverse consequences of social deprivation stress in domestic piglets. Horm Behav. 2014;65:203–210.

19. Galvão-Coelho NL, Silva HPA, De Sousa MBC. The influence of sex and relatedness on stress response in common marmosets (Callithrix jacchus). Am J Primatol. 2012;74:819–827.

20. Mendoza SP, Coe CL, Lowe EL, Levine S. The physiological response to group formation in adult male squirrel monkeys. Psychoneuroendocrinology. 1978;3:221–229.

21. Sanchez MM, McCormack KM, Howell BR. Social buffering of stress responses in nonhuman primates: Maternal regulation of the development of emotional regulatory brain circuits. Soc Neurosci. 2015;10:512–526.

22. Winslow JT, Noble PL, Lyons CK, Sterk SM, Insel TR. Rearing effects on cerebrospinal fluid oxytocin concentration and social buffering in rhesus monkeys. Neuropsychopharmacology. 2003;28:910–918.

23. Wittig RM, Crockford C, Weltring A, Langergraber KE, Deschner T, Zuberbühler K. Social support reduces stress hormone levels in wild chimpanzees across stressful events and everyday affiliations. Nature Communications. 2016;7:13361.

24. Gunnar MR, Hostinar CE. The Social Buffering of the Hypothalamic-Pituitary-Adrenocortical Axis in Humans: Developmental and Experiential Determinants. Soc Neurosci. 2015;10:479–488.

25. Ditzen B, Heinrichs M. Psychobiology of social support: the social dimension of stress buffering. Restor Neurol Neurosci. 2014;32:149–162.

26. Stanton ME, Patterson JM, Levine S. Social influences on conditioned cortisol secretion in the squirrel monkey. Psychoneuroendocrinology. 1985;10:125–134.

27. Kiyokawa Y, Hiroshima S, Takeuchi Y, Mori Y. Social buffering reduces male rats’ behavioral and corticosterone responses to a conditioned stimulus. Horm Behav. 2014;65:114–118.

28. Kiyokawa Y, Takeuchi Y, Mori Y. Two types of social buffering differentially mitigate conditioned fear responses. Eur J Neurosci. 2007;26:3606–3613.

29. Colnaghi L, Clemenza K, Groleau SE, Weiss S, Snyder AM, Lopez-Rosas M, et al. Social Involvement Modulates the Response to Novel and Adverse Life Events in Mice. PLoS ONE. 2016;11:e0163077.

30. Gunnar MR, Hostinar CE, Sanchez MM, Tottenham N, Sullivan RM. Parental buffering of fear and stress neurobiology: Reviewing parallels across rodent, monkey, and human models. Soc Neurosci. 2015;10:474–478.

31. Kingsbury L, Huang S, Wang J, Gu K, Golshani P, Wu YE, et al. Correlated Neural Activity and Encoding of Behavior across Brains of Socially Interacting Animals. Cell. 2019;178:429–446.e16.

32. Murugan M, Jang HJ, Park M, Miller EM, Cox J, Taliaferro JP, et al. Combined Social and Spatial Coding in a Descending Projection from the Prefrontal Cortex. Cell. 2017;171:1663–1677.e16.

33. Yizhar O, Fenno LE, Prigge M, Schneider F, Davidson TJ, O’Shea DJ, et al. Neocortical excitation/inhibition balance in information processing and social dysfunction. Nature. 2011;477:171–178.

34. Gilmartin MR, Balderston NL, Helmstetter FJ. Prefrontal cortical regulation of fear learning. Trends Neurosci. 2014;37:455–464.

35. Orsini CA, Kim JH, Knapska E, Maren S. Hippocampal and prefrontal projections to the basal amygdala mediate contextual regulation of fear after extinction. J Neurosci. 2011;31:17269–17277.

36. Feder A, Nestler EJ, Charney DS. Psychobiology and molecular genetics of resilience. Nat Rev Neurosci. 2009;10:446–457.

37. Maier SF, Watkins LR. Role of the medial prefrontal cortex in coping and resilience. Brain Res. 2010;1355:52–60.

38. Russo SJ, Murrough JW, Han M, Charney DS, Nestler EJ. Neurobiology of Resilience. Nat Neurosci. 2012;15:1475–1484.

39. Josselyn SA, Kohler S, Frankland PW. Finding the engram. Nat Rev Neurosci. 2015;16:521–534.

40. Denny CA, Lebois E, Ramirez S. From Engrams to Pathologies of the Brain. Frontiers in Neural Circuits. 2017;11:23.

41. Tonegawa S, Liu X, Ramirez S, Redondo R. Memory Engram Cells Have Come of Age. Neuron. 2015;87:918–931.

42. Okuyama T, Kitamura T, Roy DS, Itohara S, Tonegawa S. Ventral CA1 neurons store social memory. Science (New York, NY). 2016;353:1536–1541.

43. Sakurai K, Zhao S, Takatoh J, Rodriguez E, Lu J, Leavitt AD, et al. Capturing and Manipulating Activated Neuronal Ensembles with CANE Delineates a Hypothalamic Social-Fear Circuit. Neuron. 2016;92:739–753.

44. Bloodgood DW, Sugam JA, Holmes A, Kash TL. Fear extinction requires infralimbic cortex projections to the basolateral amygdala. Transl Psychiatry. 2018;8:60.

45. Denny CA, Kheirbek MA, Alba EL, Tanaka KF, Brachman RA, Laughman KB, et al. Hippocampal memory traces are differentially modulated by experience, time, and adult neurogenesis. Neuron. 2014;83:189–201.

46. Madisen L, Mao T, Koch H, Zhuo J, Berenyi A, Fujisawa S, et al. A toolbox of Cre-dependent optogenetic transgenic mice for light-induced activation and silencing. Nat Neurosci. 2012;15:793–802.

47. Perusini JN, Cajigas SA, Cohensedgh O, Lim SC, Pavlova IP, Donaldson ZR, et al. Optogenetic stimulation of dentate gyrus engrams restores memory in Alzheimer’s disease mice. Hippocampus. 2017;27:1110–1122.

48. Souza VR, Mendes E, Casaro M, Antiorio ATFB, Oliveira FA, Ferreira CM. Description of Ovariectomy Protocol in Mice. Methods Mol Biol. 2019;1916:303–309.

49. Drew MR, Denny CA, Hen R. Arrest of adult hippocampal neurogenesis in mice impairs single-but not multiple-trial contextual fear conditioning. Behav Neurosci. 2010;124:446–454.

50. Sparta DR, Stamatakis AM, Phillips JL, Hovelsø N, van Zessen R, Stuber GD. Construction of implantable optical fibers for long-term optogenetic manipulation of neural circuits. Nat Protoc. 2011;7:12–23.

51. Stamatakis AM, Stuber GD. Activation of lateral habenula inputs to the ventral midbrain promotes behavioral avoidance. Nat Neurosci. 2012;15:1105–1107.

52. Kim Y, Venkataraju KU, Pradhan K, Mende C, Taranda J, Turaga SC, et al. Mapping social behavior-induced brain activation at cellular resolution in the mouse. Cell Rep. 2015;10:292–305.

53. Barker JM, Taylor JR, Chandler LJ. A unifying model of the role of the infralimbic cortex in extinction and habits. Learn Mem. 2014;21:441–448.

54. Thompson BM, Baratta MV, Biedenkapp JC, Rudy JW, Watkins LR, Maier SF. Activation of the infralimbic cortex in a fear context enhances extinction learning. Learn Mem. 2010;17:591–599.

55. Lungwitz EA, Stuber GD, Johnson PL, Dietrich AD, Schartz N, Hanrahan B, et al. The role of the medial prefrontal cortex in regulating social familiarity-induced anxiolysis. Neuropsychopharmacology. 2014;39:1009–1019.

56. Hinwood M, Tynan RJ, Day TA, Walker FR. Repeated social defeat selectively increases deltaFosB expression and histone H3 acetylation in the infralimbic medial prefrontal cortex. Cerebral Cortex (New York, NY: 1991). 2011;21:262–271.

57. Lehmann ML, Herkenham M. Environmental enrichment confers stress resiliency to social defeat through an infralimbic cortex-dependent neuroanatomical pathway. The Journal of Neuroscience: The Official Journal of the Society for Neuroscience. 2011;31:6159–6173.

58. Matochik JA. Role of the main olfactory system in recognition between individual spiny mice. Physiology & Behavior. 1988;42:217–222.

59. Okuyama T. Social memory engram in the hippocampus. Neurosci Res. 2018;129:17–23.

60. Phillips ML, Robinson HA, Pozzo-Miller L. Ventral hippocampal projections to the medial prefrontal cortex regulate social memory. Elife. 2019;8.

61. Kitamura T, Sun C, Martin J, Kitch LJ, Schnitzer MJ, Tonegawa S. Entorhinal Cortical Ocean Cells Encode Specific Contexts and Drive Context-Specific Fear Memory. Neuron. 2015;87:1317–1331.

62. Liu X, Ramirez S, Pang PT, Puryear CB, Govindarajan A, Deisseroth K, et al. Optogenetic stimulation of a hippocampal engram activates fear memory recall. Nature. 2012;484:381–385.

63. Ramirez S, Liu X, MacDonald CJ, Moffa A, Zhou J, Redondo RL, et al. Activating positive memory engrams suppresses depression-like behaviour. Nature. 2015;522:335–339.

64. Redondo RL, Kim J, Arons AL, Ramirez S, Liu X, Tonegawa S. Bidirectional switch of the valence associated with a hippocampal contextual memory engram. Nature. 2014;513:426–430.

65. Cai DJ, Aharoni D, Shuman T, Shobe J, Biane J, Song W, et al. A shared neural ensemble links distinct contextual memories encoded close in time. Nature. 2016;534:115–118.

66. Lacagnina AF, Brockway ET, Crovetti CR, Shue F, McCarty MJ, Sattler KP, et al. Distinct hippocampal engrams control extinction and relapse of fear memory. Nat Neurosci. 2019;22:753–761.

67. Chen BK, Murawski NJ, Cincotta C, McKissick O, Finkelstein A, Hamidi AB, et al. Artificially Enhancing and Suppressing Hippocampus-Mediated Memories. Curr Biol. 2019;29:1885–1894.e4.

68. Adhikari A, Lerner TN, Finkelstein J, Pak S, Jennings JH, Davidson TJ, et al. Basomedial amygdala mediates top-down control of anxiety and fear. Nature. 2015;527:179–185.

69. Guzman YF, Tronson NC, Sato K, Mesic I, Guedea AL, Nishimori K, et al. Role of oxytocin receptors in modulation of fear by social memory. Psychopharmacology (Berl). 2014;231:2097–2105.

70. Sabihi S, Dong SM, Maurer SD, Post C, Leuner B. Oxytocin in the medial prefrontal cortex attenuates anxiety: Anatomical and receptor specificity and mechanism of action. Neuropharmacology. 2017;125:1–12.

71. Kiyokawa Y, Hennessy MB. Comparative studies of social buffering: A consideration of approaches, terminology, and pitfalls. Neurosci Biobehav Rev. 2018;86:131–141.

72. Kiyokawa Y, Takeuchi Y. Social buffering ameliorates conditioned fear responses in the presence of an auditory conditioned stimulus. Physiol Behav. 2017;168:34–40.

73. Hall BS, Romeo RD. The influence of poststress social factors on hormonal reactivity in prepubertal male rats. Dev Psychobiol. 2014;56:1061–1069.

74. Romeo RD. Perspectives on stress resilience and adolescent neurobehavioral function. Neurobiol Stress. 2015;1:128–133.

75. Sterley T-L, Baimoukhametova D, Füzesi T, Zurek AA, Daviu N, Rasiah NP, et al. Social transmission and buffering of synaptic changes after stress. Nat Neurosci. 2018;21:393–403.

76. Burkett JP, Andari E, Johnson ZV, Curry DC, de Waal FB, Young LJ. Oxytocin-dependent consolation behavior in rodents. Science (New York, NY). 2016;351:375–378.

77. Guzman YF, Tronson NC, Jovasevic V, Sato K, Guedea AL, Mizukami H, et al. Fear-enhancing effects of septal oxytocin receptors. Nat Neurosci. 2013;16:1185–1187.

78. Kiyokawa Y, Honda A, Takeuchi Y, Mori Y. A familiar conspecific is more effective than an unfamiliar conspecific for social buffering of conditioned fear responses in male rats. Behav Brain Res. 2014;267:189–193.

79. Kiyokawa Y, Kikusui T, Takeuchi Y, Mori Y. Partner’s stress status influences social buffering effects in rats. Behav Neurosci. 2004;118:798–804.

80. Tottenham N. Social scaffolding of human amygdala-mPFCcircuit development. Soc Neurosci. 2015;10:489–499.

