## Supplemental figures for "Optogenetic reactivation of prefrontal social memory trace mimics social buffering of fear"

**Supplementary figures**

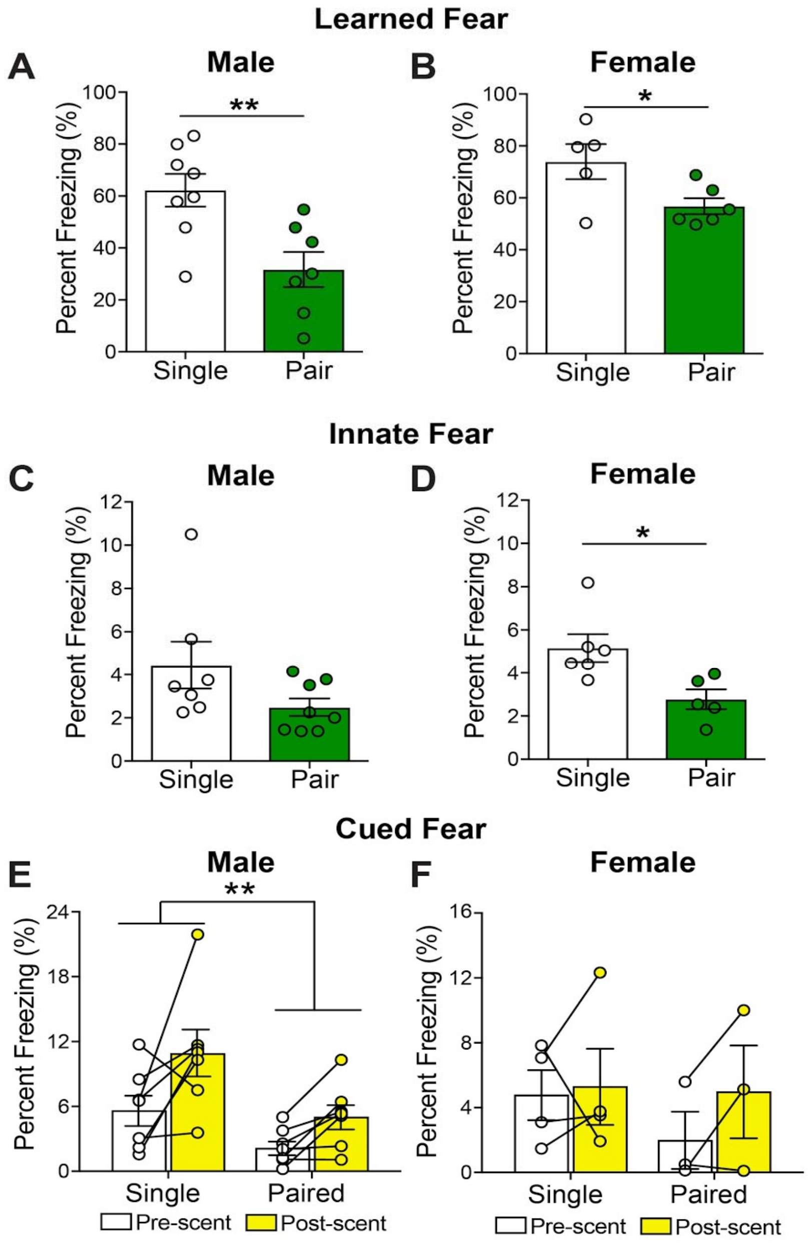


**Figure S1.** Male and female mice exhibit social buffering. (A-B) Results from Figure 1E separated by male and female experimental mice. Male mice exhibit overall higher levels of freezing, but both males and females show reduced freezing when a cagemate is present. (C-D) Cagemate presence also reduced freezing in a novel aversive environment, although this was significant only for females. (E-F) Males displayed a more robust response to fear cue (lemon scent) and stronger socially-mediated reduction in freezing than females. Error bars represent ± SEM. * $p < 0.05$; ** $p < 0.01$, *** $p < 0.001$.

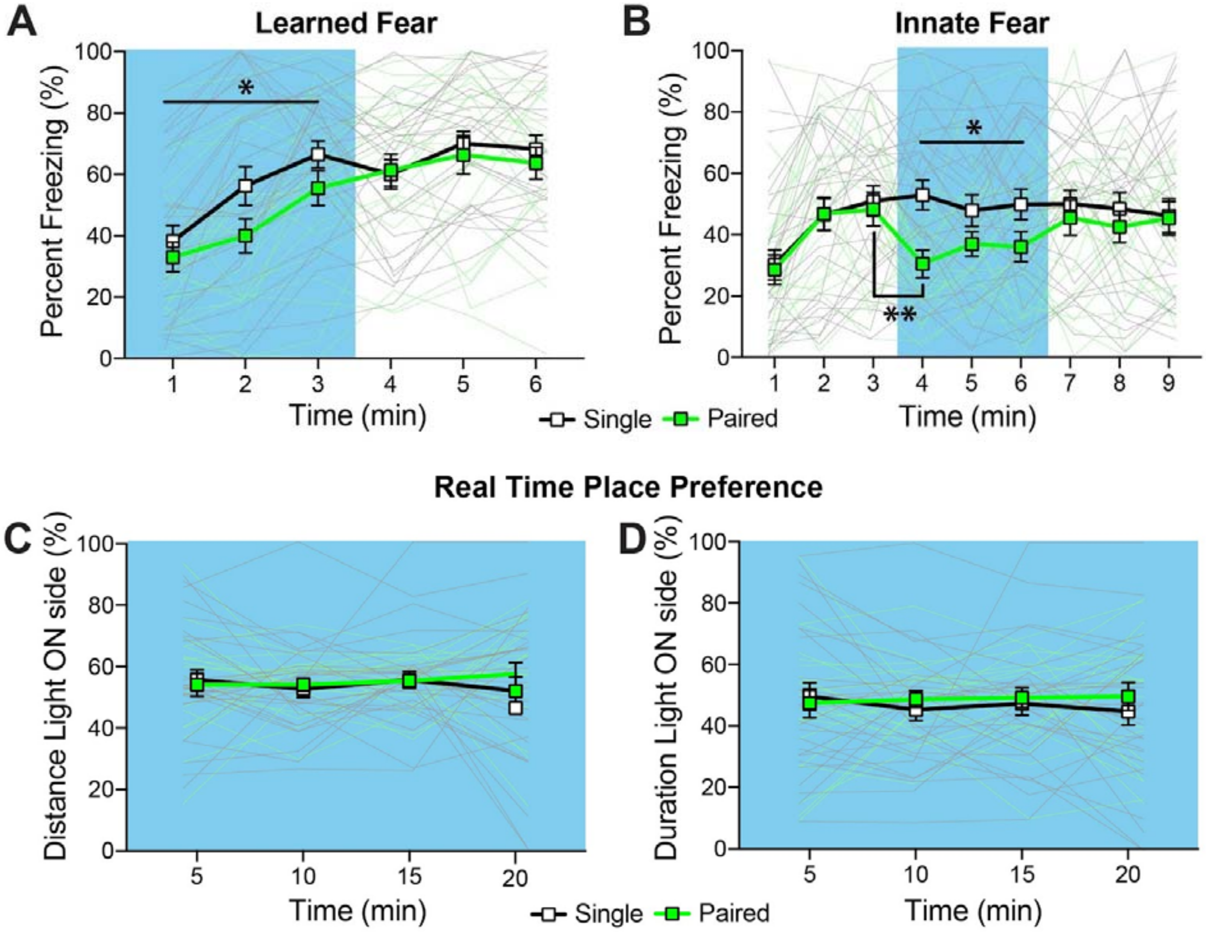


**Figure S2.** Optogenetic re-activation of the social memory trace reduces expression of learned and innate fears, but is not rewarding. (A) Optogenetic reactivation of socially-labeled ensembles in the IL-PFC caused a reduction in freezing during exposure to a previously shock-paired context compared to controls. (B) Optogenetic reactivation of socially-labeled ensembles significantly reduced freezing in a novel context during light-on period compared to controls in a. (C-D) No differences in distance traveled or duration in the light-on side of the place preference chamber were observed during reactivation of socially-labeled ensembles in the IL-PFC in a 20 minute test. Blue = light ON. Error bars represent ± SEM. * $p < 0.05$; ** $p < 0.01$, *** $p < 0.001$.
